# Precise loading of scarce reagents on droplet microarrays

**DOI:** 10.64898/2026.03.17.712446

**Authors:** Shenghao Tan, Jon E. Albo, Nate J. Cira

## Abstract

Many experiments rely on expensive or scarce liquids, such as costly reagents, or biological samples available only in limited quantities. Droplet microarrays are an especially promising approach to conserving these materials because they support highly parallelized reactions in small volumes. However, existing droplet microarray loading methods based on discontinuous dewetting suffer from loading inconsistencies and large dead volumes. In this work, we present the Small Volume Loader (SVL) for the Surface Patterned Omniphobic Tiles (SPOTs) platform that enables precise deposition on droplet microarrays while minimizing reagent waste. By establishing a physical model of the loading process, we identified that deposition volume is governed by the sum of hydrostatic and Laplace pressures at the reservoir outlet. To optimize performance, we engineered a pressure-compensating flared reservoir geometry that maintains constant total pressure regardless of the remaining liquid level. This design ensures that the deposited volume is independent of reservoir volume and reduces dead volume to 5 μL. We demonstrated the platform’s utility through high-throughput elicitor screening for natural antimicrobial production from *Streptomyces venezuelae*. The resulting assays used 100-fold less material than conventional methods, allowing us to conduct over 32,000 assays with modest quantities of starting material. This enabled us to identify specific stressors that optimize the production of the antibiotics chloramphenicol and jadomycin B. Together, we demonstrated improved loading performance for droplet microarray platforms, allowing precise, accessible, and high-throughput assays using only minimal volumes of scarce materials.

## Introduction

Limited availability of laboratory reagents due to their cost (e.g., antibodies) or scarcity (e.g., patient samples, synthesized compounds, natural products) presents a challenge in many research settings. Droplet microarrays enable thousands of parallelized experiments, each confined in individual picoliter-to microliter-scale droplets containing drugs, reagents, and cells^1–3^. These arrays greatly reduce reagent consumption while improving experimental throughput in molecular detection and quantification, combinatorial screening, and reaction optimization^4–7^. The precise partitioning of liquid reagents on a droplet microarray is critical for accurate and repeatable experimental results.

Multiple approaches have been used to deposit liquids on droplet microarrays. Microinjector and piezoelectric-based droplet printing methods^6–10^ offer active control over the droplet volume; however, they often require translational stages and precise actuation mechanisms that add cost and are limited in throughput. Passive loading methods such as droplet-dragging^5,11,12^, dipping^13–16^, shearing^17^, and flooding^18,19^ rely on discontinuous dewetting to rapidly partition and deposit onto physically or chemically patterned surfaces. These strategies however are prone to the volume variation influenced by sliding speed, sliding angle, reagent volume, and viscosity or surface tension of the bulk liquid^15,19^.

We previously introduced the Surface Patterned Omniphobic Tiles (SPOTs) platform and demonstrated its versatility in different applications^20,21^. The SPOTs platform consists of two components: 1) a glass plate with circular -philic area (spots) created by laser ablation, surrounded by a superomniphobic background 2) superomniphobic loaders that retain and partition liquids. We identified the gap height, defined by the distance between the loader and the plate, as a crucial factor for the deposition volume. While the original loader designs were efficient and precise for loading common reagents, they required a minimum reservoir volume (dead volume) far greater than the amount ultimately deposited onto the SPOTs plates, making them unsuitable for handling scarce reagents. Furthermore, the physical basis of the relationship between the deposition volume and the spot size remained unexplored.

Here we present an improved loading device for droplet microarrays, offering minimal dead volume and improved loading consistency. To achieve this, we first introduced a physical model that predicts volume deposition. We then enhanced the loader performance by modifying the reservoir geometry. To demonstrate the loader performance, we performed a highly parallelized antimicrobial metabolite screening and phenotypic characterization for metabolite elicitation from *Streptomyces venezuelae*.

## Results and discussion

### SVL assembly achieves consistent volume deposition

In earlier work, we implemented “slot” loaders to dispense a single reagent type across an entire SPOTs plate, and “row” loaders to distribute multiple reagent types across many rows or columns on one axis, similar to filling a microplate with multi-channel micropipettes. However, we observed that different liquid volumes in the reservoir can change the deposition volume. To assess the effect of the volume remaining in the reservoir on the volume deposited on the spots, we continuously dispensed 250 μL of fluorescein solution across three SPOTs plates with a “row” loader and quantified droplet volumes via spectrophotometry. Interestingly, the deposition volume exhibited a non-monotonic dependence on the remaining liquid volume (Fig. 1b, blue line). As the liquid level dropped, the deposition volume decreased gradually at first. However, once the remaining volume fell below 35 μL, the deposition volume rose sharply despite continued depletion. In addition, “slot” loaders and “row” loaders have relatively high dead volume, below which loading becomes inconsistent or fails.

**Figure 1.**
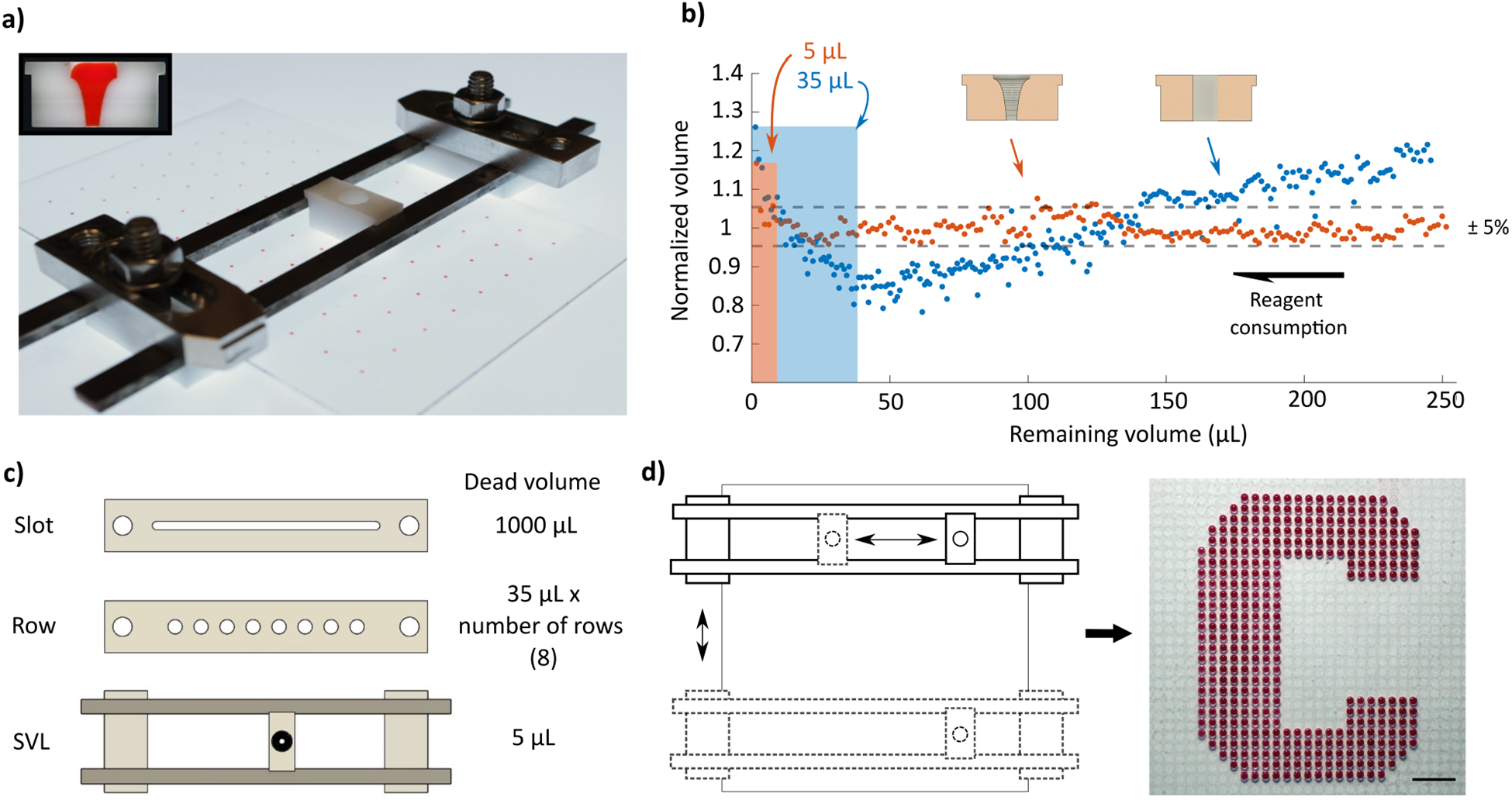
Overview of the small volume loader (SVL). (a) A SVL assembly deposits droplets onto a SPOT array, with minimal dead volume, by featuring a flared cross-section. (b) Comparison of deposition volume profiles between SVL (flared) geometry and the original geometry with a 250 μL initial volume in the reservoir. The cylindrical reservoir exhibits a non-monotonic loading profile, while the SVL reservoir maintains a uniform deposition volume independent of the volume remaining in the reservoir. The colored areas represent the corresponding dead volume of each loader. (c) Unique reservoir geometry enables minimal dead volume in SVL compared to previous loader iterations (d) The multi-axial motions of the SVL offer flexible spatial control, allowing users to generate custom microdroplet patterns, scale bar = 10 mm.

To resolve the non-monotonic loading trend and reduce dead volumes, we introduce a loader configuration termed Small Volume Loader (SVL) assembly (Fig. 1a). The assembly comprises plastic loaders to retain and partition reagents and metal rails to support loader movements. The SVL assembly allows for two-axis movement across the SPOTs plate, facilitating customized and complete utilization of the reagents, as well as selective patterning (Fig. 1d). We further modified the reservoir geometry to achieve uniform deposition with a single loading of up to 250 μL (Fig. 1b orange line). The SVL assembly demonstrated enhanced loading consistency compared to earlier loader iterations (Fig. S1) and significantly reduced dead volume by 200x (compared to 1000 μL in “slot” loaders) and 7x (compared to 35 μL in “row” loaders) (Fig. 1c), while being capable of row/column and entire-plate modes of loading.

### Pressure-dependent droplet model predicts deposition volume

To elucidate the trend in deposition volume, we first sought to understand the physics of loading. We measured the loaded volume *V*_*l*_ as a function of the diameter of laser-ablated spots, *D*_*s*_ (Fig. 2a). When the spot diameter is larger than the gap height (distance between the bottom of the loader and top of the plate),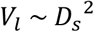, consistent with a disc-like droplet, with its height restricted by the gap height H_g_ between the loader and the SPOTs plate. At smaller diameters, the relationship deviates from this trend, motivating a deeper consideration of the loading physics.

**Figure 2.**
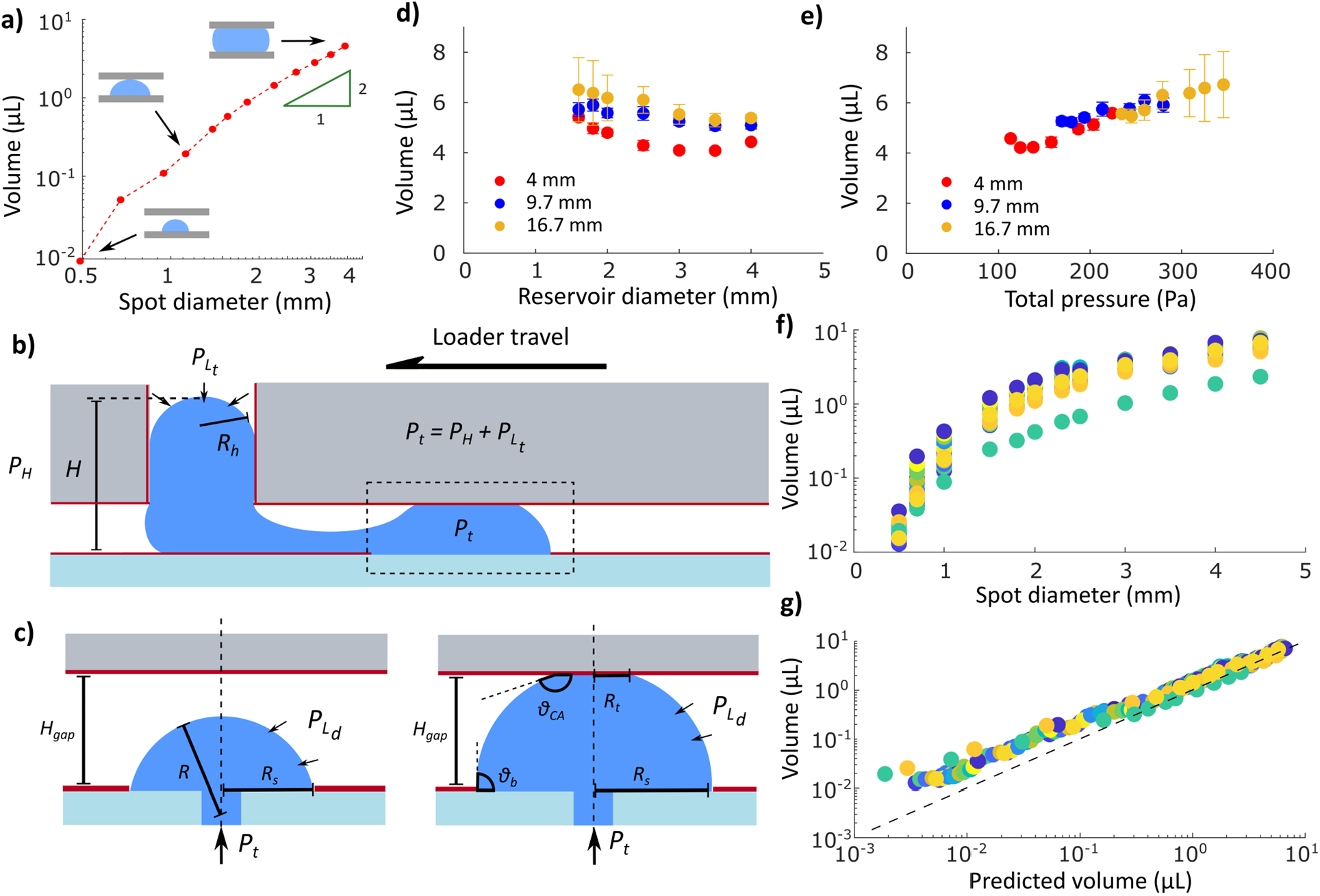
Deposition volume is dependent on pressure in the liquid at the spot. (a) Spot diameter and deposition volume relationship with a “slot” loader. (b) A cross-sectional view of the loading process. The Laplace pressure and hydrostatic pressure combine to set a total pressure, which influences the droplet shape (schematic not drawn to scale). (c) Proposed simplified model to predict the shape and volume of a droplet from an imposed total pressure P_t_. (Left) A spherical cap prediction is used until the droplet contacts the bottom of the loader, then a squeezed droplet prediction is used (Right). (d) Reservoir diameter and loader height both independently influence deposition volume (e) Experimental volumes collapse into a single curve when plotted against the total pressure, suggesting that total pressure influences deposited volume of the droplet, regardless of the pressure source. (f) Spot size vs. deposition profile of 24 loader configurations with various heights, reservoir diameter, and gap heights (Supplementary Table 1). (g) Measured deposition volume compared with predicted volume calculated with the total pressure-controlled squeezed droplet model.

To understand the deposition mechanism, we monitored the wetting and dewetting process during loading as the loader passes over a spot (Supplementary Video 1). The spot is immediately wet upon coming into contact with the liquid in the reservoir, forming a liquid bridge between the loader outlet and the spot. As the loader moves over and beyond the spot, the liquid bridge is extended, and eventually, as the loader outlet moves away from the spot, the liquid bridge breaks, leaving a volume of liquid deposited on the spot. The droplet shape on the spot must resist the liquid pressure coming from the reservoir immediately before liquid bridge rupture. We consider the process before breakup to be pseudo-static, due to the low capillary number of the loading process ^21^. We hypothesized that the deposited volume is strongly related to the liquid geometry on the spot immediately before the liquid bridge breaks.

The reservoir outlet and the spot being loaded are at the same vertical height, hence should have the same pressure. Therefore, we next examine the pressure contributions from the reservoir. The total pressure at the bottom of the reservoir has contributions from two sources, Laplace and hydrostatic pressure. When the liquid is dispensed into the cylindrical reservoir with a superomniphobic surface, it forms a cylindrical column capped by a spherical-cap meniscus at the top of the loading reservoir (Fig. 2b). The spherical-cap meniscus maintains a consistent radius of curvature at different reservoir volumes, due to the coating’s low sliding angle, resulting in a consistent top Laplace pressure 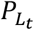. As the liquid is consumed, the column height decreases, thereby reducing the hydrostatic pressure *P*_*H*_. Based on these observations, we hypothesize that *V*_*l*_ depends on the total pressure at the reservoir outlet, *P*_*t*_ = *P*_*H*_ + *P*_*Lt*_ . To test our hypothesis, we fabricated loaders with varying heights and reservoir diameters and measured the corresponding *V*_*l*_. Consistent with this picture, we found that increasing the loader height and reducing the reservoir diameter both increase deposition volume for a given spot size (Fig. 2d).

When we consider the total pressure (sum of Laplace pressure and hydrostatic pressure), the deposited volumes across different loader heights and different loader reservoir sizes collapse to a single curve, consistent with the total pressure being the key driver of the deposition volume, independent of the pressure source (Fig. 2e). Furthermore, higher *P*_*t*_ values increased the variability of *V*_*l*_, likely due to fluctuations in the volume partition dynamics during the pinch-off event (Supplementary Video 1).

Following this understanding that the total pressure imposed by the loader determines the deposited volume, we constructed a simplified, quasi-static droplet model to predict the deposition volume. In this model, we determine which droplet shape has the correct curvature to generate a droplet Laplace pressure 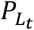, that equates to the pressure in the liquid at the bottom of the reservoir, *P*_*t*_.

To simplify the complex geometry of a droplet with liquid bridge connected laterally to the reservoir, we instead consider an axisymmetric approximation without the liquid bridge, as if droplet were filled by injection through the substrate by a pressure source *P*_*t*_ onto a spot of radius *R*_*s*_. The droplet is pinned at the edge of the spot due to the surface energy difference between the exposed glass surface and the surrounding superomniphobic background. Under low *P*_*t*_ and small *R*_*s*_, the droplet forms a spherical cap with the diameter *R*_*s*_, where the volume is given by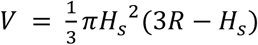, with radius of curvature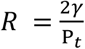, and the cap height 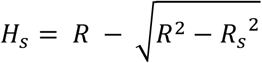(Fig. 2c). At higher *P*_*t*_ and larger *R*_*s*_, the droplet becomes tall enough to touch the bottom of the loader and becomes “squeezed” (Fig. 2d). Due to the superomniphobicity of the loader surface, the droplet always maintains a large contact angle with minimal hysteresis at the top surface. In this regime, we determine the droplet shape by solving the axisymmetric form of the Young-Laplace equation: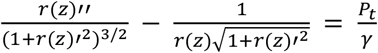 with the initial conditions *r*(0) = *R*_*s*_ and *r*(0) = −*cot*(*θ*_*b*_), where *r*(*z*) is the droplet radius as a function of the height *z*. We omit the gravitational contribution to this shape due to the small change in hydrostatic pressure across the narrow gap height, which then dictates that the droplet surface has constant mean curvature, forming the slice of an unduloid^22^. The droplet profile is numerically solved by iteratively increasing the base contact angle *θ*_*b*_ until the condition *r*(*H*_*gap*_)’ = *cot*(θ) is satisfied, where *θ*_*CA*_ is the contact angle of the liquid on superomniphobic surface and *H*_*gap*_ is the gap height. To validate the model, we fabricated loaders with different *H, H*_*gap*_, and *R*_*h*_ (Table S1), resulting in variable loading volumes (Fig. 2g), and found good agreement between the predictions and the measurements (Fig. 2h).

Notably, this model does not include the volumetric contribution of the liquid bridge and pinch-off dynamics. Deposition volume is somewhat underestimated by this simple model for smaller spots, where the spherical cap approximation is applied, suggesting that the liquid bridge contributes a larger fraction of the volume under these conditions (Supplementary material Vid 1). In practice, empirically calibrated fits are used for experiments to ensure loading accuracy. Nonetheless, the model helps establish total volume *P*_*t*_ as the key factor for the loading process and inspires strategies to maintain consistent deposition volume.

### Pressure balance by loader geometry ensures consistent deposition volume

The physical model allows us to consider the implications of different loader reservoir geometries. For the earlier straight-walled reservoirs, the depositing droplet experiences fluctuating pressure, first from the decreasing hydrostatic pressure when liquid column reduces in height as the reagents are continuously partitioned onto spots. Second, as the liquid in the reservoir is further depleted, the deposition volume changes markedly once the bulk liquid detaches from the reservoir wall and forms a spherical shape on the highly non-wetting surfaces. At this stage, *P*_*t*_ increases rapidly as the reduction in hydrostatic pressure is surpassed by the rise in Laplace pressure. By calculating the total pressure with different amounts of reagent volume in cylindrical reservoirs, we found that the decrease and increase in deposition volume follows the calculated pressure changes (purple line, fig. 3a).

**Figure 3.**
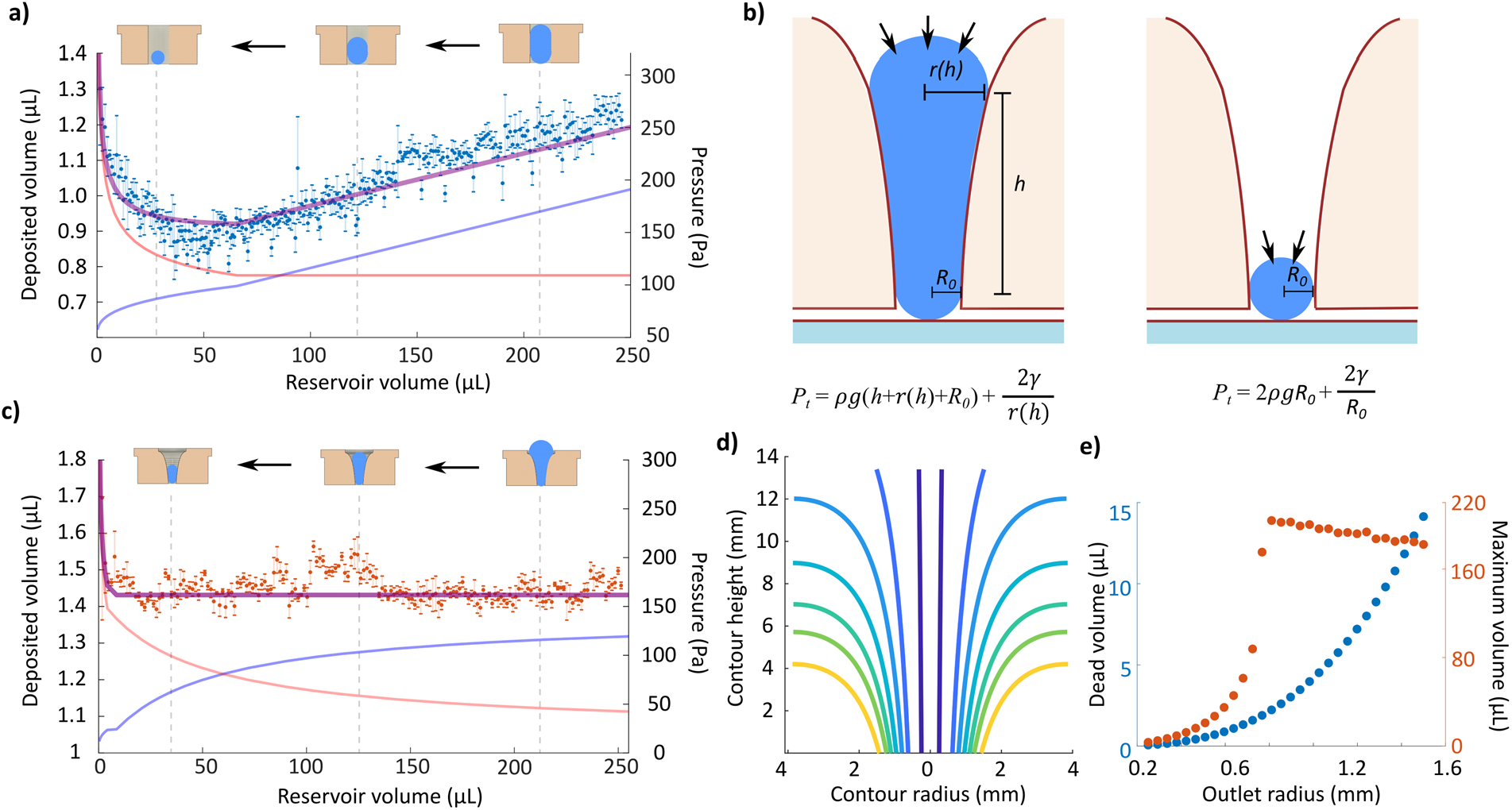
Flared reservoir geometry ensures loading consistency by pressure compensation. (a) Pressure-volume profile of a straight-hole reservoir. As the liquid volume reduces, changes in hydrostatic pressure (blue line) and Laplace pressure (red line) result in non-monotonic total pressure (purple line) and loading profile (blue markers). (b) Physical basis of flared reservoir geometry. In order to achieve constant pressure, independent of the height of the liquid in the loader, the total pressure during the loading process (hydrostatic pressure from liquid column and spherical cap, plus Laplace pressure from the spherical cap, left) must equate to the total pressure at the dead volume (hydrostatic and Laplace pressure from the spherical cap, right). (c) Pressure-volume profile of a flared reservoir. The total pressure (purple line) remains constant throughout the loading process, ensuring consistent deposition volume (red markers) until the dead volume is reached. (d) Flared reservoir contours calculated for different outlet diameters. (e) Comparing the dead volume to maximal volume of flared reservoir loaders with various outlet sizes. The total reservoir height is capped when *r*(*h*) *→ ∞*, up to 13.4 mm, based on the loader material stock.

To minimize the pressure variation, we introduced a flared reservoir geometry, reasoning that as the reservoir volume decreases, the corresponding drop in hydrostatic pressure can be compensated by an increase in Laplace pressure from the reservoir contour *r*(*h*), thereby maintaining a consistent total pressure *P*_*t*_ throughout the loading process. Specifically, *r*(*h*) is defined by the relationship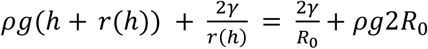 where *R*_0_ is the radius of the reservoir outlet and *r*(*h*) > *R*_0_ . The reservoir geometry is given by

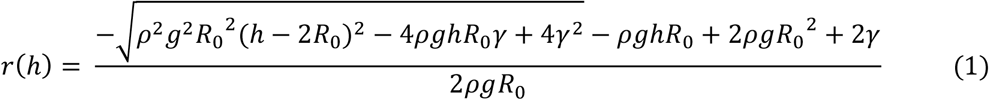

and ensures that *P*_*t*_ is consistent until the dead volume is reached (fig. 3b). As the reservoir volume decreases, the corresponding drop in hydrostatic pressure is compensated by an increase in Laplace pressure from the reservoir contour *r*(*h*), thereby maintaining a consistent total pressure *P*_*t*_ and consistent deposition volume throughout the loading process. We note that the calculation assumes the radius of curvature of the spherical cap matches the local reservoir radius *r*(*h*); however, deviations may arise at larger flaring angles. Deformation due to gravity is also neglected. Nevertheless, we found that the loading consistency through the full reservoir volume is greatly improved (Fig. 3c).

Consequently, the dead volume is determined by the outlet size rather than by the flaring reservoir geometry. Minimizing *R*_0_ would be desirable to achieve minimal dead volume; however, the total height of the loader needs to be drastically increased to achieve a reasonable reservoir capacity (Fig. 3d), making fabrication and manipulation difficult. To balance dead volume and reservoir capacity, we designed Small Volume Loaders with a 1 mm outlet radius, achieving consistent loading with a total of 250 μL of liquid and a dead volume of 5 μL (Fig. 3c, Fig. 3e).

To determine whether loading consistency is influenced by reagent properties, we examined the volume deposition dynamics using a fluorescein solution containing two concentrations of ethanol. Due to the lower surface tension of these liquids, under the same reservoir geometry, the top Laplace pressure reduces; however, the curvature also increases to generate sufficient droplet Laplace pressure, rendering higher deposition volume for the same total pressure. As expected, we observed a gradual decrease in deposition volume as the reservoir volume decreased, caused by an insufficient reduction in reservoir radius with decreasing liquid height (Fig. S3). To mitigate the volume variation, the reservoir geometry can be easily modified according to Eq. 1 to accommodate liquids with different surface tension and density, or repeated loading cycles can be performed (Fig. S3). Consistent deposition volumes were obtained across different loading speeds, with only a moderate increase in variability at excessively high traversal speeds (Fig. S4).

### High-throughput elicitor screening for antimicrobial metabolite production

Natural products play prominent roles in medicine; however, they are difficult to obtain in large quantities, especially at the screening stage. This challenge is especially critical for antibiotics, which are vital to medicine, but have become increasingly difficult to discover. Prominent producers of medically important antibiotics, such as members of the genus *Streptomyces*, often produce antibiotics in limited quantities and under specific culture media compositions^23,24^, environmental stress factors^25,26^, and microbial interactions^27^. Therefore, optimizing culture conditions for maximal production or selective secretion of specific antibiotics (elicitation) demands large-scale efforts. In addition, conventional screening strategies, such as bioactivity assays performed in well plates, often require large quantities of metabolites obtained from bulk-scale fermentations and subsequent labor-intensive liquid extractions.

Although numerous microfluidic approaches^28–31^ have been implemented for high-throughput *Streptomyces* cultivation and antimicrobial evaluation, these methods generally provide only binary phenotypic outcomes (kill / no kill) and lack quantitative information on production rate, metabolite concentration, and metabolite identity/purity. Moreover, *Streptomyces* metabolomes are typically variable and chemically complex^32–34^, complicating the identification of bioactive compounds and necessitating dereplication strategies^35–37^.

To address the shortcomings in existing screening platforms, we applied our improved droplet microarray system to high-throughput elicitor screening for antimicrobial metabolite production. We selected *Streptomyces venezuelae* ISP 5230 as a model producer, as it produces two chemically distinct antimicrobial metabolites in two different media, chloramphenicol in maltose-yeast extract-malt extract (MYM) and jadomycin B^38,39^ in glucose-isoleucine (GI) medium^38^. To elicit and improve antimicrobial metabolite production, we supplemented the base media with elicitors including antibiotics, stress factors, microbial extracts, and signaling molecules. Cultures were incubated for 24, 48, and 72 hours, after which the extracted metabolites were distributed across 108 spots. To assess if elicitor addition altered or expanded antibiotic production profiles, we evaluated the phenotypic susceptibility of three target organisms: *Escherichia coli, Bacillus subtilis* wild type, and *Bacillus subtilis* chloramphenicol-resistant (cm^R^), against each extract. We assessed the growth of each organism at different dilutions of each extract, to elucidate the concentration and identity of the produced antimicrobial metabolites.

SPOTs plates with uniform spot dimensions were loaded with LB broth inoculated with each target strain and overlaid with dried metabolite plates to generate 1 μL assays incorporating a 12-step, 1.4-fold dilution series. By measuring which metabolite dilutions caused inhibition, we can quantify the antibiotic production for each elicitor across each day. Accounting for different targets, serial dilutions, replicates, media, elicitors, and multiple timepoints, the full experiment involved 32,400 individual assays with 900 tests performed per SPOTs plate, giving us detailed information regarding antibiotic production in each media formulation. The use of small volume loaders reduced the quantity of metabolites required for each antimicrobial sensitivity test (AST) by approximately 100-fold compared to conventional 96-well microtiter plate assays (∼1 μL on SPOTs vs ∼100 μL in 96-well format). Consequently, we were able to culture the producers in 24-well plates to assess the metabolite production from each elicitor condition and each time point, as opposed to arrays of larger flasks. In comparison, if ASTs were conducted in conventional 96-well plates, this screening would have required 28.8 L of producer culture and 338 96-well plates. The enhanced maneuverability of the small-volume loaders enabled greater spot density compared to previous “row” loaders (900 spots vs. 384 spots per plate) ^21^, while still allowing precise, discrete reagent deposition across rows. The minimum inhibitory concentration (MIC_50_) values were determined based on microbial growth after 12 hours, quantified via dark-field imaging (Fig. 4b).

**Figure 4.**
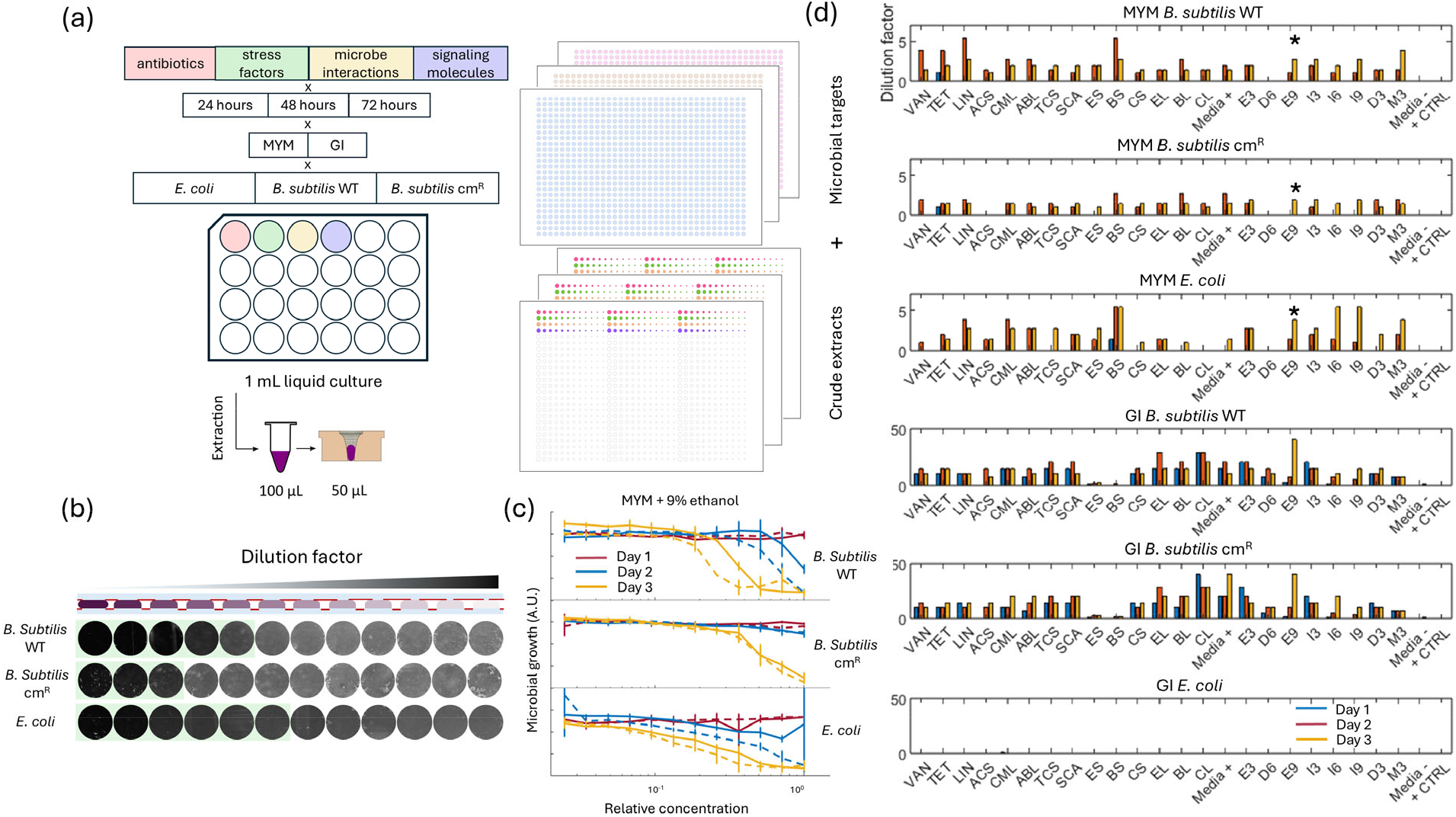
High-throughput elicitor screening and antimicrobial metabolite characterization. (a) Antibiotic production, extraction, and assay workflow. 1 mL *S. venezuelae* cultures are exposed to a diverse panel of potential elicitors, and the extractions are performed after 24, 48, and 72 hours to obtain 100 μL crude extracts, and deposited onto SPOTs plates with SVLs. Each plate is sandwiched to another SPOTs plate with LB broth inoculated with one of the target bacterial strains, to obtain the minimum inhibitory dilution factors representing the concentration of potency of the produced antibiotics (b) Darkfield images of target bacteria growth exposed to the crude extracts at different dilutions, t = 16 hours. (c) Representative susceptibility profiles of target bacteria strains. The half-max inhibition dilution factor is determined as the dilution at which less than 50% of bacteria growth occurred compared to positive controls. (d) Half-max inhibitory dilution factor against the target strains of crude extracts produced from *S. venezuelae* cultures with various elicitors at 24, 48, and 72 hours (see table S1 for abbreviations). Star signs indicate data derived from (c).

The platform performed robustly, revealing diverse production profiles across the culture conditions. Replicates within each plate showed consistent results, indicating reliable volume deposition across varying reservoir volumes (Fig. 4c). We obtained detailed inhibition responses from each target strain against each crude extract (Fig. S6) and compressed the data to display antibiotics production dynamics in each condition (Fig. 4d). *B. subtilis* cm^R^ exhibited higher resistance against metabolites from organisms grown in MYM than the wild type strain, suggesting predominant chloramphenicol production, as expected. Extracts from GI cultures displayed little to no activity against *E. coli*, implying predominant jadomycin B production with minimal chloramphenicol biosynthesis, since *E. coli* is more resistant to jadomycin B compared to *B. subtilis* (Fig. S4). In general, jadomycin B accumulated rapidly, reaching peak levels within 24 hours, whereas chloramphenicol production occurs gradually across 72 hours.

Interestingly, supplementation of GI media with 10% *E. coli* and *B. subtilis* spent media drastically suppressed jadomycin B production, and occasionally elicited chloramphenicol production, unanticipated in the GI media background (Fig. S6). The most pronounced enhancement of chloramphenicol yield occurred with the addition of lincomycin, *B. subtilis* spent media, and alcohols (Fig. 4d), with modest improvements observed for other antibiotic elicitors, consistent with previously reported elicitation effects in *Streptomyces spp*^40–42^. Conversely, only the addition of 9% ethanol significantly enhanced jadomycin B production, albeit with a reduced production rate. Overall, we achieved ∼2.5x chloramphenicol and ∼2x jadomycin B production relative to the standard production media without the addition of elicitors.

## Conclusion

In this work, we presented an improved loader assembly for the SPOTs platform, developed from a detailed understanding of the loading physics. By introducing a flared reservoir geometry to maintain constant total pressure, SVLs ensure consistent deposition volume with minimal dead volume. The same principles can be applied to improve the performance of “slot” and “row” loaders. Leveraging this enhanced liquid-handling capability, we established a high-throughput workflow for elicitor screening and susceptibility profiling aimed at enhancing antimicrobial metabolite production. The platform addresses two major limitations of conventional screening, high metabolite consumption and low throughput, by drastically reducing assay volumes while maintaining precision. Our experiments demonstrate the utility of this platform for precise manipulation of scarce reagents in natural product discovery, microbiology, and molecular biology applications.

## Acknowledgements

We thank K. Gallagher for providing the spore stock of *Streptomyces venezuelae* ISP 5230. We thank B. Kirby for helpful discussion. Some microbial strains used in this work were provided by the USDA-ARS Culture Collection (NRRL). Research reported in this publication was supported by the National Institute of General Medical Sciences (NIGMS) of the National Institutes of Health under award number: R35GM157104. The content is solely the responsibility of the authors and does not necessarily represent the official views of the National Institutes of Health.

## Methods

### Fabrication of small volume loaders

The loader pieces were designed in Fusion 360 (Autodesk) and prototyped in Delrin acetal homopolymer (McMaster Carr, GA, US) with a Nomad 883 desktop CNC machine (Carbide 3D, US). Scaled injection molding production was performed by Respon Manufacturing (Shenzhen, China). The loader rail was assembled with precision ground 4410 steel bars (McMaster Carr), CNC milled Delrin rail legs, and bolts and washers for assembly.

### Volume measurements and prediction

The deposition volume of SVLs on SPOTs plates were measured empirically via spectrophotometry. The different sized spots on a SPOTs plate were loaded with 24 g/L fluorescein disodium by selected loaders. The SPOTs plate was interfaced with a 96-well plate (Corning) loaded with 200 μL of deionized water and inverted repeatedly to mix. The absorptions at 490, 520, and 530 nm were taken on a SpectraMax 384 spectrophotometer (Molecular Devices) and compared to established standard curves^21^. Volume estimations based on the squeezed droplet model were performed with a custom MATLAB program.

Briefly, the program initially assumes a small bottom contact angle *θ*_*b*_ and numerically calculates the droplet contour based on Young-Laplace equation from the total pressure *P*_*t*_. The top contact angle *θ*_*t*_ at the loader surface is calculated from the contour and compared to the droplet contact angle on a superomniphobic surface *θ*_*CA*_ = 175°. *θ*_*b*_ is increased or decreased iteratively, with incrementally smaller angular steps, until |*θ*_*CA*_ *− θ*_*t*_| < 0.1°.

### Culture, extraction, and evaluation of antimicrobial crude extracts

*Streptomyces venezuelae* ISP5230 culture was inoculated from spore stock in MYM media (4 g/L maltose, 4 g/L yeast extract, 10 g/L malt extract) supplemented with 1.9 g/L MOPS for 12 hours at 30 °C. The Glucose-Isoleucine (GI) media were prepared in accordance with Jakeman et al^43^. The MSM solution was supplemented with 60 mM isoleucine, 33 mM glucose, and 0.05 mM phosphate ^43^. *S. venezuelae* was inoculated in MYM media and GI media at 0.8 and 0.005 OD600, respectively. Elicitor concentrations were determined by minimum inhibitory concentration (MIC) assays so that *S. venezuelae* growth wasn’t inhibited, and the negative control extracts do not inhibit *E. coli, B. subtilis* WT and *B. subtilis* cm^R^. The selected elicitor concentrations are 0.04 μg/mL vancomycin, 0.08 μg/mL tetracycline, 2 μg/mL lincomycin, 10 μg/mL acetylglucosamine, 0.3125 μg/mL chloramphenicol, 10 μg/mL α-Amino-γ-butyrolactone, 0.0025 μg/mL triclosan, 2 μg/mL scandium chloride, 10% *E. coli, B. subtilis* and *Candida albicans* supernatants and lysates (24 hour culture in LB, LB, Sabouraud broth, respectively), 3% and 6% DMSO, 3%, 6%, 9% isopropanol, 3% methanol, 3%, 9% ethanol. In the MYM production medium, alcohols were added as specified. In the GI medium, elicitor alcohols replaced ethanol as the shock agent, while all other elicitors were supplemented with 3% ethanol. Each condition was established in 1 mL liquid culture in 24-well plates and incubated at 30 °C @ 250 rpm.

The secondary metabolites were extracted after 24, 48, and 72 hours of incubation with 1 mL of ethyl acetate. 1 mL of ethyl acetate was added to each inoculum; the mixture is vortexed vigorously, briefly centrifuged, and the ethyl acetate layer was carefully extracted. The solvent was evaporated under vacuum, and the crude extracts were resuspended in 100 μL of DI water.

### Minimal volume metabolite AST fingerprinting

50 μL of each resuspended crude extract was pipetted into the small volume loader piece and distributed across 108 spots with various sizes. The droplets were allowed to dry to ensure consistent final volume. Another SPOTs plate with uniform 2.5 mm spots and matching layout was loaded with one of the target organisms (*E. coli, B. subtilis WT, B. subtilis* cm^R^) inoculated in LB broth at 0.005 OD and sandwiched with the crude extract plates. The SPOTs plate assemblies were incubated at 37 °C for 16 hours and imaged with an automated microscope (Nikon Eclipse Ti-2) under a darkfield optical configuration.

