## Supplementary Information for "Precise loading of scarce reagents on droplet microarrays"

**Supplementary Materials**

**Supplementary video 1:** This video shows the loading dynamics from below, looking through the substrate plate. First the liquid wets the spot, then, as the loader moves past the spot, a liquid bridge forms and the liquid pinches off from the loader.

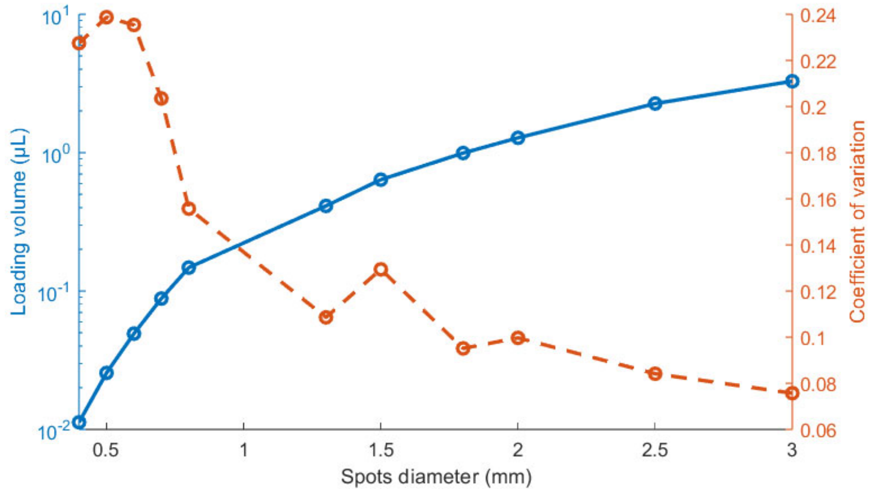


**Figure S1 | Spots diameter and deposition volume relationship with SVL on SPOTs plates.** The deposition volume variation is comparable to earlier loader iterations[21].

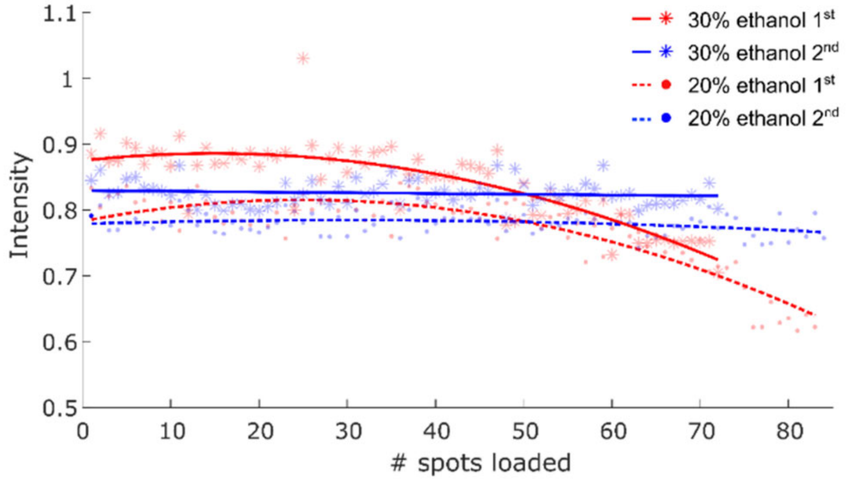


**Figure S2 | Influence of surface tension on deposition volume.** Fluorescein solutions supplemented with 20% and 30% ethanol deposited less volume on the substrate as the reservoir volume decreases (red). Repeated loading renders volume consistent across spots (blue).

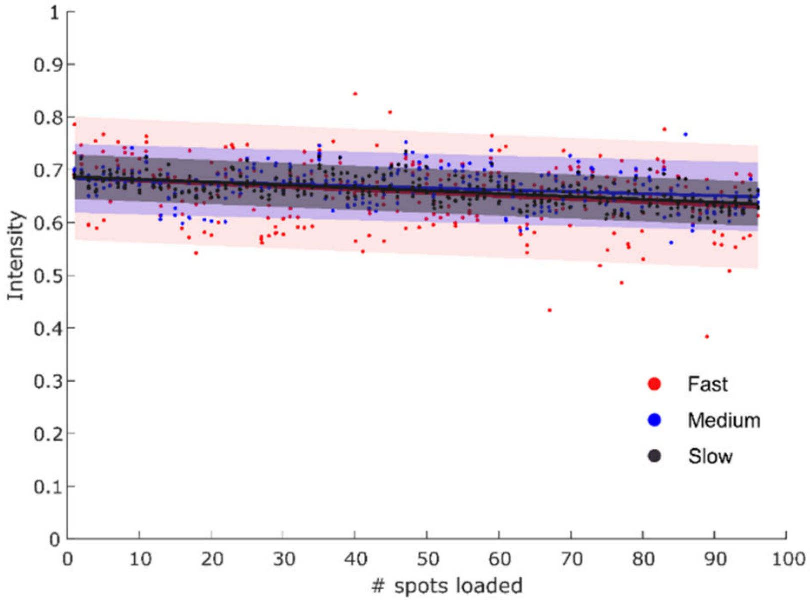


**Figure S3 | Influence of loading velocity on deposition volume.** Colored shadings represent 95% confidence interval of a linear fit.

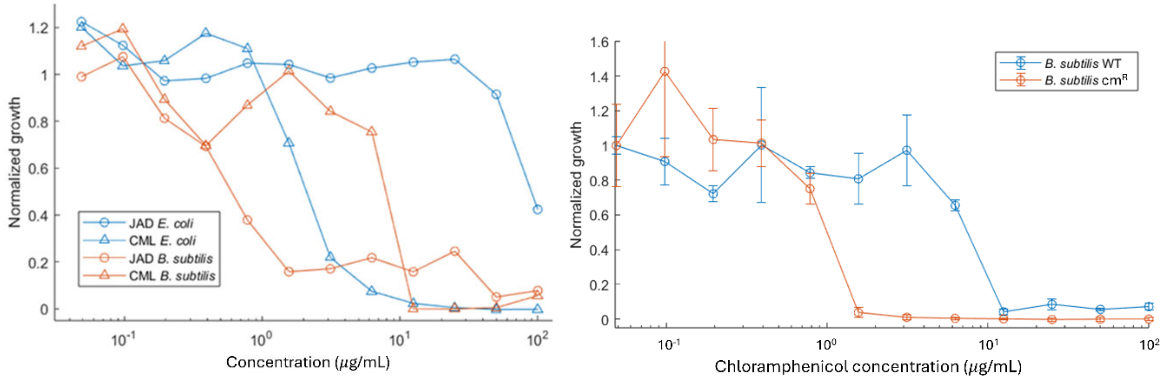


**Figure S4 | Nominal MIC profiles of *E. coli*, *B. subtilis* WT, and *B. subtilis* cm$^{R}$ against chloramphenicol and jadomycin B.** These MIC assays are performed in 96 well-plates.

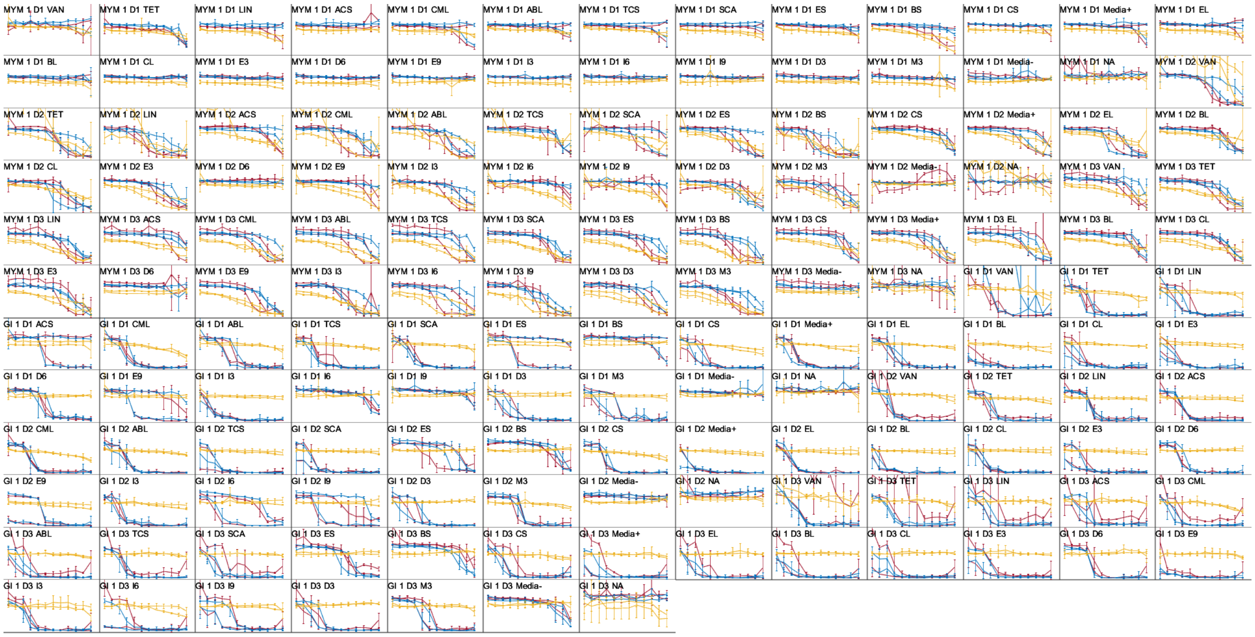


**Fig. S5 | Complete metabolite data.** Each curve is derived from 3 replicates of a 12-spot dilution series with the metabolites derived from 1 mL of producer culture. Same colored lines represent replicates of producer culture from different wells, from which the metabolites are extracted separately. Error bars represent the standard deviation of three replicate assays performed at each concentration for each producer culture. X-axis: relative concentration. Plot titles indicate background media, day on which producer culture was extracted, and elicitor (see table S1 for abbreviation key). Y-axis: normalized growth of *E. coli* (yellow lines), *B. subtilis* WT (red lines), *B. subtilis* $cm^R$ (blue lines).

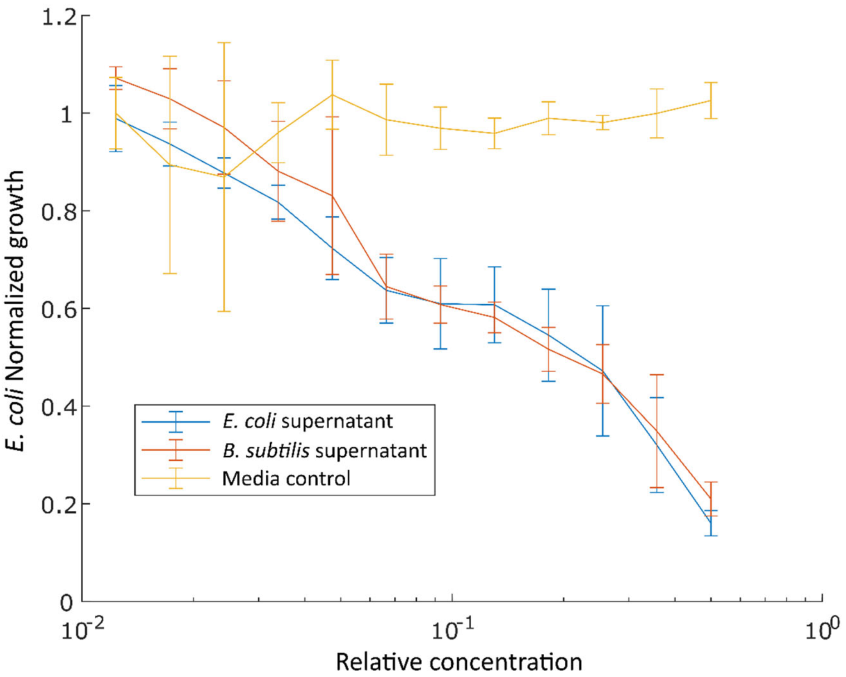


**Fig. S6 | Chloramphenicol elicitation from *S. venezuelae* in GI medium supplemented with spent media from *E. coli* and *B. subtilis*.** Crude extracts from *S. venezuelae* cultures supplemented with 10% *E. coli* and *B. subtilis* spent media displayed inhibition against *E. coli*, which is highly resistant to jadomycin B. This suggests the production of chloramphenicol, which is not expected from this producer in a GI medium.

| Elicitor | Abbreviation |
|---|---|
| vancomycin | VAN |
| tetracycline | TET |
| lincomycin | LIN |
| acetylglucosamine | ACS |
| chloramphenicol | CML |
| amino-butyrolactone | ABL |
| triclosan | TCS |
| scandium (III) chloride | SCA |
| *E. coli* supernatant | ES |
| *B. subtilis* supernatant | BS |
| *C. albicans* supernatant | CS |
| *E. coli* lysate | EL |
| *B. subtilis* lysate | BL |
| *C. albicans* lysate | CL |
| ethanol 3% | E3 |
| DMSO 6% | D6 |
| ethanol 9% | E9 |
| isopropanol 3% | I3 |
| isopropanol 6% | I6 |
| isopropanol 9% | I9 |
| DMSO 3% | D3 |
| methanol 3% | M3 |

**Table S1 | Elicitor abbreviations.**
